# Remodeling of *Mycobacterium tuberculosis* lipids regulates *prpCD* during acid growth arrest

**DOI:** 10.1101/509570

**Authors:** Jacob J. Baker, Robert B. Abramovitch

## Abstract

*Mycobacterium tuberculosis* (Mtb) establishes a state of non-replicating persistence when it is cultured at acidic pH with glycerol as a sole carbon source. Growth can be restored by spontaneous mutations in the *ppe51* gene or supplementation with pyruvate, supporting that acid growth arrests is a genetically controlled, adaptive process and not simply a physiological limitation associated with acidic pH. Transcriptional profiling identified the methylcitrate synthase and methylcitrate dehydratase genes (*prpC* and *prpD*, respectively) as being selectively induced during acid growth arrest. *prpCD* along with isocitrate lyase (*icl*) enable Mtb to detoxify propionyl-CoA through the methylcitrate cycle. The goal of this study was to examine mechanisms underlying the regulation of *prpCD* during acid growth arrest. Induction of *prpCD* during acid growth arrest was reduced when the medium was supplemented with vitamin B12 (which enables an alternative propionate detoxification pathway) and enhanced in an *icl* mutant (which is required for the propionate detoxification), suggesting that Mtb is responding to elevated levels of propionyl-CoA during acidic growth arrest. We hypothesized that an endogenous source of propionyl-CoA generated during metabolism of methyl-branched lipids may be regulating *prpCD*. Using Mtb radiolabeled with ^14^C-propionate or ^14^C-acetate, it was observed that lipids are remodeled during acid growth arrest, with triacylglycerol being catabolized and sulfolipid and trehalose dimycolate being synthesized. Blocking TAG lipolysis using the lipase inhibitor tetrahydrolipstatin, resulted in enhanced *prpC* induction during acid growth arrest, suggesting that lipid remodeling may function, in part, to detoxify propionate. Notably, *prpC* was not induced during acid growth arrest when using lactate instead of glycerol. We propose that metabolism of glycerol at acidic pH may result in the accumulation of propionyl-CoA and that lipid remodeling may function as a detoxification mechanism.

**Importance:** During infection, *Mycobacterium tuberculosis* (Mtb) colonizes acidic environments, such as the macrophage phagosome and granuloma. Understanding regulatory and metabolic adaptations that occur in response to acidic pH can provide insights int0 mechanisms used by the bacterium to adapt to the host. We have previously shown that Mtb exhibits pH-dependent metabolic adaptations and requires anaplerotic enzymes, such as Icl1/2 and PckA, to grow optimally at acidic pH. Additionally, we have observed that Mtb can only grow on specific carbon sources at acidic pH. Together these findings show that Mtb integrates environmental pH and carbon source to regulate its metabolism. In this study, it is shown that Mtb remodels its lipids and modulates the expression of propionyl-CoA detoxifying genes *prpCD* when grown on glycerol at acidic pH. This finding suggests that lipid remodeling at acidic pH may contribute to detoxification of propionyl-CoA, by incorporating the metabolite into methyl-branched cell envelope lipids.

## Introduction

Survival of *Mycobacterium tuberculosis* (Mtb) during infection requires sensing and adapting to the diverse and adverse environments of the host. Growing evidence suggests that the mechanisms of Mtb adaptation during infection are unique from those required during *in vitro* culture. First, many genes dispensable for growth *in vitro* have been shown to be essential during infection(1, 2). Similarly, past efforts that identified antimycobacterial drugs with potent *in vitro* activity have been unable to achieve *in vivo* efficacy, possibly due to the distinct *in vivo* environment and the physiological state of Mtb in response to that environment(3). Thus, development of effective therapy for Mtb requires careful consideration of the environmental conditions encountered by Mtb during infection.

A growing body of research supports that adaptation to acidic pH is important to Mtb pathogenesis. Mtb colonizes mildly acidic environments (~pH 6.5-5.0) in the macrophage phagosome and induces genes regulated by the acidic pH-inducible PhoPR two component regulator(4–6). Mutants that are defective in maintaining cytosolic pH-homeostasis (*e.g. marP*(7, 8)) or altering gene expression in response to acidic pH (*phoPR(5)*) are strongly attenuated for growth in macrophages and infected mice(5, 7, 9), supporting that acid tolerance and pH-dependent adaptations are required for Mtb pathogenesis.

Mtb growth at acidic pH in minimal medium requires host-associated carbon sources that function at the intersection of glycolysis and the TCA cycle (*e.g*. pyruvate, acetate, oxaloacetate and cholesterol)(10). In contrast, in other tested carbon sources (such as glucose or glycerol), Mtb fully arrests its growth at acidic pH, establishes a state of non-replicating persistence (NRP), and becomes tolerant to antibiotics(11). Growth-arrested Mtb is resuscitated by the addition of pyruvate showing that growth arrest is due to a pH-dependent checkpoint on metabolism or nutrient acquisition. Additionally, mutations in PPE51 enable Mtb to grow in acidic medium with glycerol as a sole carbon source, demonstrating that acid growth arrest is a regulated process. This acidic pH- and carbon source-dependent NRP is a new model of Mtb persistence, that is referred to as acid growth arrest(11), and provides new opportunities to conduct studies examining mechanisms underlying how pH regulates Mtb growth, metabolism, persistence and transcriptional networks.

Transcriptional profiling identified several genes with strong induction at acidic pH, independent of carbon source, including genes associated with anaplerotic metabolism (*e.g. icl1, pckA, ppdK*, and *mez*), respiration (*e.g*. type 2 NADH dehydrogenase, *ndh*), and amino acid/nitrogen metabolism (*e.g*. arginine biosynthesis, *argCDFGHR*). *Icl1/2* and *pckA* are induced at acidic pH in both carbon sources and we have previously shown that they are required for optimal growth and metabolic remodeling at acidic pH(11). In addition to inducing genes involved in the anaplerotic node, we also observed that Mtb induces a subset of genes specifically during acidic pH growth arrest(10). Notably, these genes were not induced at acidic pH when supplemented with the growth permissive carbon source pyruvate. Among these acidic pH growth arrest-induced genes were two genes involved in the methylcitrate cycle, encoding methylcitrate synthase and methylcitrate dehydratase (*prpC* and *prpD*, respectively). *prpCD*, along with *icl1* which is also a methylisocitrate lyase, have been characterized for their role in the detoxification of propionyl-CoA intermediates generated from the catabolism of cholesterol as well as branched- and odd-chain fatty acids (12–16). The *prpCD* operon is induced by cholesterol, propionate and hypoxia and this induction is dependent on the SigE-regulated Rv1129c regulator (16, 17). However, Mtb cultured at acidic pH in a defined minimal medium does not have an exogenous source of propionyl-CoA, cholesterol or other branched chain precursors. Given the lack of an obvious propionyl-CoA source in the minimal medium conditions of acidic pH growth arrest, understanding the mechanisms of induction of *prpCD* at acidic pH could provide further insights into the metabolic state of Mtb at acidic pH. Mtb remodels its lipids at acidic pH, including the induction of sulfolipid, 2,3-diacyltrehaloses (DAT) and penta-acyltrehaloses (PAT)(10). These lipids and other long-chain fatty acids contain methyl-branched lipids, which are preferentially labelled when Mtb is provided ^14^C-propionate(18, 19). The goal of this study was to test the hypothesis that *prpCD* is induced during acid growth arrest due to presence of endogenous propionyl-CoA released during lipid remodeling.

## Results

### Transcriptional induction of prpCD during acidic growth arrest

Induction of *prpCD* at acidic pH in minimal medium with glycerol as a sole carbon source was verified by quantitative PCR (Figure 1A). We hypothesized that the observed *prpCD* induction is the result of increased endogenous production of propionyl-CoA during acidic pH growth arrest. Previous work has shown that supplementation of vitamin B12 is able to relieve the requirement for prpCD-mediated propionyl-CoA metabolism by opening the alternative methylmalonyl pathway (14). The supplementation of vitamin B12 significantly reduced *prpCD* expression at both pH 7.0 and pH 5.7 (Figure 1B). Similarly, in the Δ*icl1/2* mutant that lacks methylisocitrate lyase activity, *prpC* expression was increased ~2-fold at pH 5.7 as compared to WT Mtb (Figure 1C). Together, these results support the proposal that *prpCD* induction at acidic pH is linked to propionyl-CoA metabolism.

**Figure 1.**
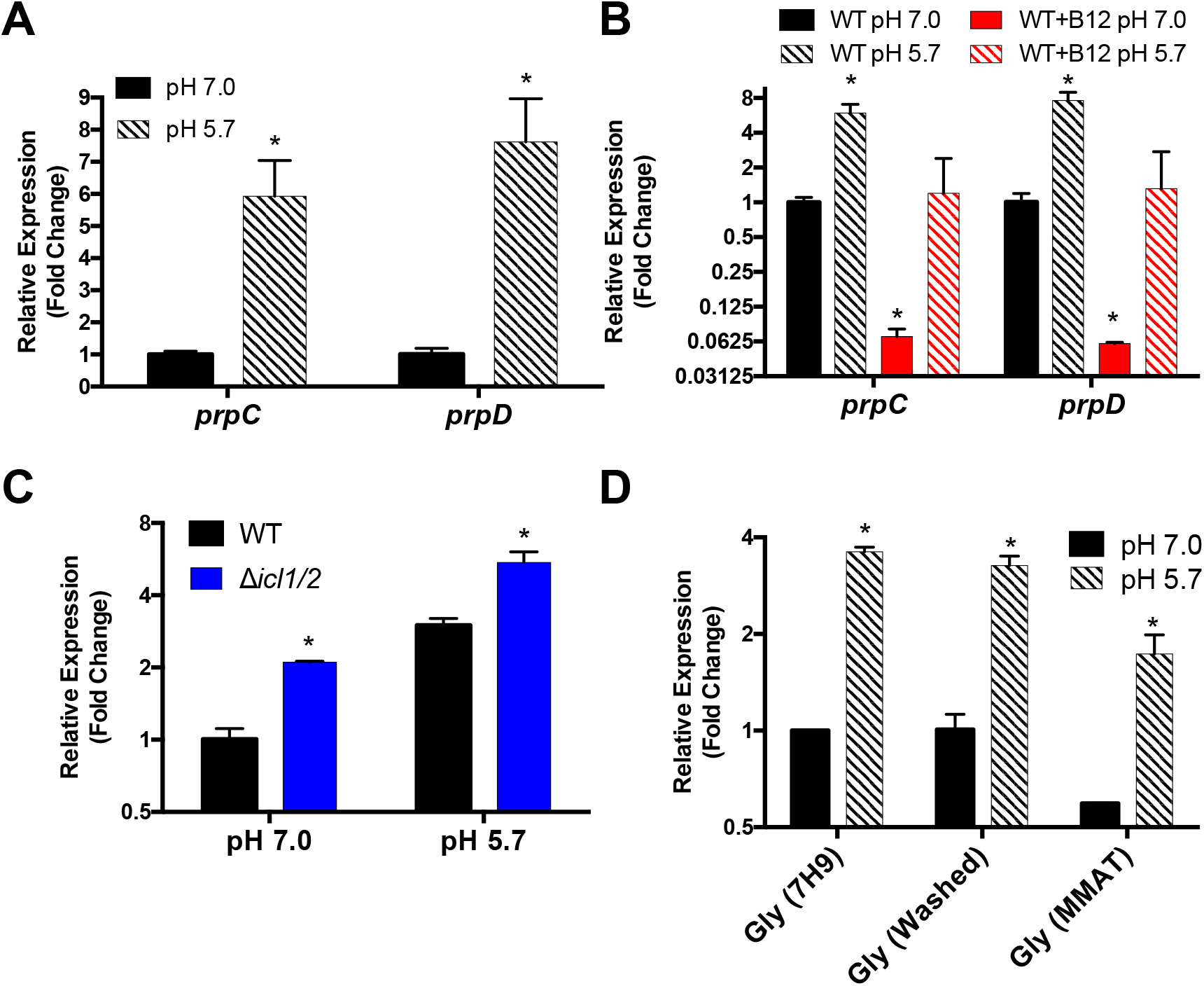
*prpCD* is induced at acidic pH and responds to alterations in propionyl-CoA metabolism. A) Quantitative real time PCR (qPCR) of *prpC* and *prpD* mRNA at pH 7.0 and pH 5.7 with glycerol as a single carbon source confirms that *prpCD* are induced during acidic pH growth arrest. * Significantly induced (*p*<0.05) at pH 5.7 relative to pH 7.0 (t test). B) *prpCD* expression at pH 7.0 and pH 5.7 is reduced with addition of vitamin B12. * Significantly induced (*p*<0.05) at pH 5.7 relative to pH 7.0 (t test). C) *prpC* expression at pH 7.0 and pH 5.7 is increased in the *Δicl/12* mutant. * Significantly induced (*p*<0.05) at in *icl1/2* mutant relative to WT (t test). D) *prpC* induction at pH 5.7 in minimal medium with glycerol as a single carbon is observed in Mtb that was maintained prior to the experiment in 7H9+OADC rich medium-(Gly [7H9]), in 7H9+OADC and washed 3 times in PBS + 0.05% Tween 80 (Gly [Washed]), and in Mtb cultured in minimal medium with glycerol as a single carbon source buffered to pH 7.0 (Gly [MMAT]) * Significantly induced (*p*<0.05) at pH 5.7 relative to pH 7.0 (t test). All experiments repeated at least twice with two independent biological replicates.

Because *prpCD* expression at acidic pH responds to changes in propionyl-CoA metabolism, we sought to identify the source of propionyl-CoA that could lead to *prpCD* induction. The rich medium 7H9+OADC may contain some propionyl-CoA sources from the supplemented albumin, therefore, we tested whether there was carryover of these carbon sources into the minimal medium culture. After washing Mtb cultures 3 times in minimal medium prior to transfer to acid growth arrest medium, *prpC* was still induced following 3 days of acidic growth arrest (Figure 1D). Furthermore, Mtb grown from a frozen stock grown exclusively in minimal medium containing glycerol as a single carbon source still induced *prpC* at pH 5.7 with glycerol as a sole carbon source (Figure 1D). Together, these results suggest that the induction of *prpCD* is not due to an exogenous propionyl-CoA source.

### Mtb cell envelope remodeling under acidic pH growth arrest modulates prpCD induction

Given the absence of an exogenous propionyl-CoA source, we hypothesized that one source of propionate during pH 5.7 growth arrest could be the breakdown of Mtb cell envelope or storage lipids. To test this hypothesis, Mtb was cultured in rich medium in the presence of ^14^C-propionate or ^14^C-acetate for 3 weeks in order to radiolabel Mtb lipids. This radiolabeled Mtb was then inoculated into minimal medium at pH 5.7 containing glycerol as a single carbon source and the relative abundance of lipid species was measured over time during pH 5.7 growth arrest. The total radioactivity of the samples decreased by less than 10% through the 12-day time course (Figure 2A); however, over the same time period the relative abundance of radiolabeled triacylglycerol (TAG) decreased to one-fourth of the initial concentration while the relative abundance of both trehalose dimycolate (TDM) and sulfolipid (SL) increased ~4-fold (Figure 2B-H). This result suggests that during pH 5.7 growth arrest, Mtb may utilize endogenous TAG to remodel its cell wall through the increased synthesis of both TDM and SL.

**Figure 2.**
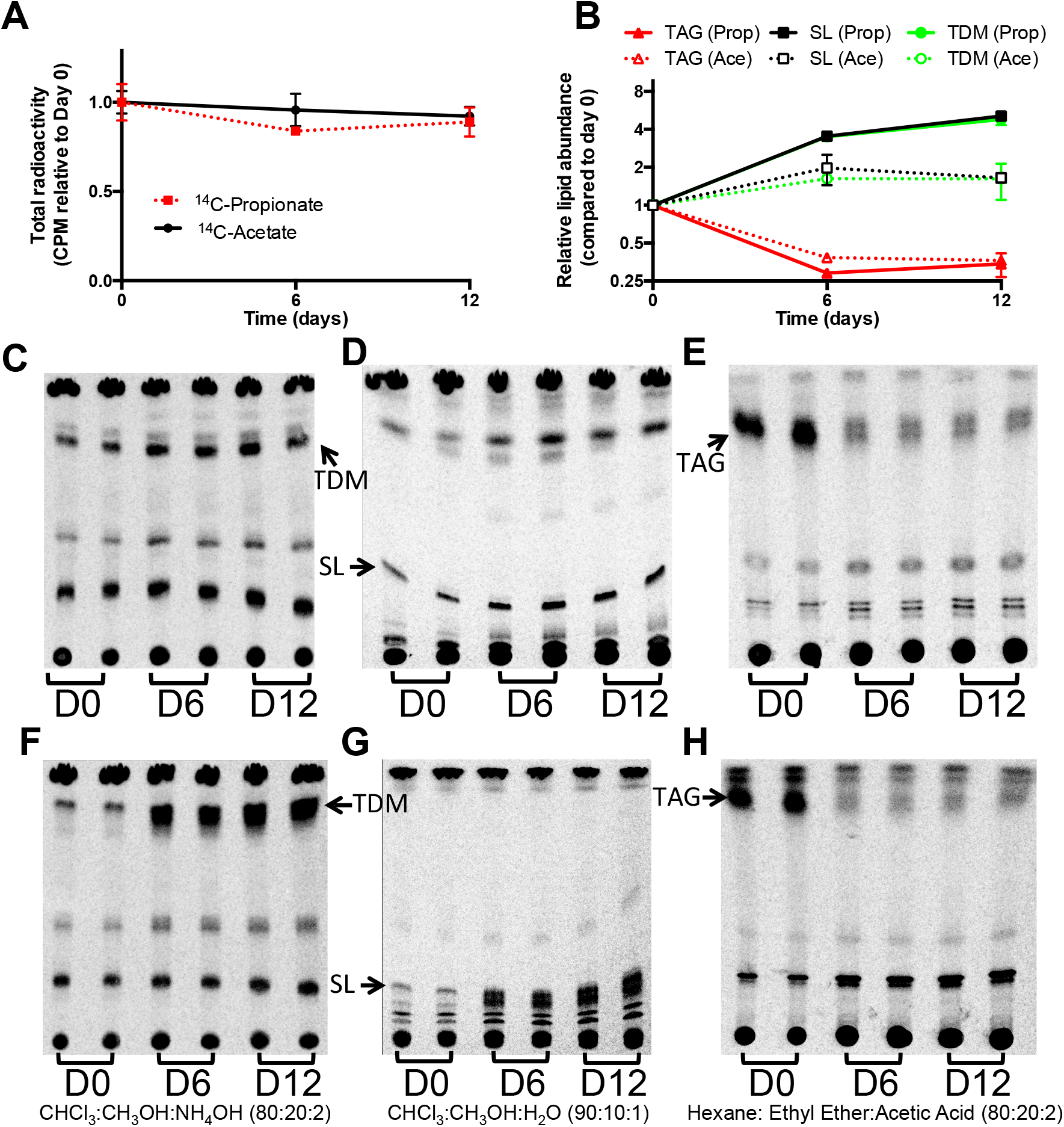
Mtb utilizes endogenous TAG for the synthesis of TDM and SL at acidic pH. Mtb was grown in the presence of ^14^C-acetate or ^14^C-propionate for 3 weeks prior to transferring to minimal medium containing glycerol as a single carbon source buffered to pH 5.7. A) Total radioactivity of Mtb whole cells over time. Over 12 days a ~10% reduction in radioactivity was observed. B) Relative lipid species abundance of triacylglycerol (TAG), sulfolipid (SL), and trehalose dimycolate (TDM) over time in Mtb labelled with ^14^C-acetate (Ace) or ^14^C-propionate (Prop). C-H) Thin Layer Chromatography (TLC) images showing relative abundance of TAG, SL and TDM at 0, 6, and 12 days after transfer of ^14^C-acetate-(C-E) or ^14^C-propionate- (F-H) labeled Mtb to acidic pH growth arrest (D0, D6, and D12, respectively).

To test the hypothesis that lipid remodeling is a source of endogenous propionyl-CoA during pH 5.7 growth arrest, we sought to disrupt the ability of Mtb to metabolize TAG to SL and TDM. The addition of the lipase inhibitor tetrahydrolipstatin (THL) to Mtb cultures blocked the remodeling of Mtb TAG to SL and TDM (Figure 3A-C). Interestingly, despite blocking lipid remodeling, treatment with THL increased *prpC* expression during pH 5.7 growth arrest 3-fold compared to DMSO treated Mtb (Figure 3D). This result suggests that lipid remodeling of TAG to SL and TDM is not a source of *prpCD* induction at acidic pH; instead, the increase in *prpC* induction with addition of THL suggests that lipid remodeling at acidic pH may act as a mechanism to relieve propionyl-CoA stress.

**Figure 3.**
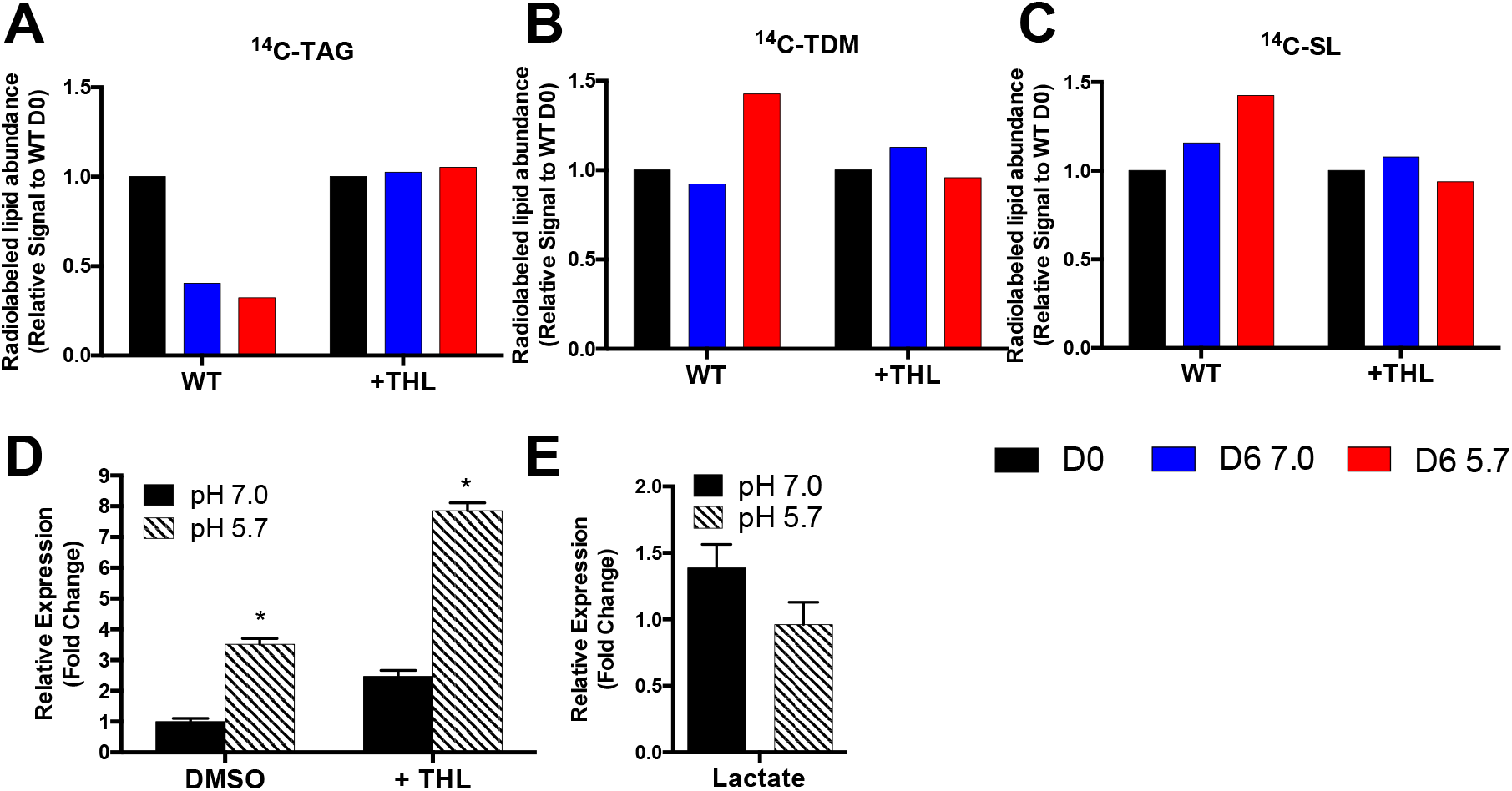
Inhibition of lipid remodeling at acidic pH increases *prpC* induction. A-C) Remodeling of radiolabeled TAG, TDM, and SL at day 0 (D0) and after incubation for 6 days at pH 7.0 or pH 5.7 (D6 7.0 or D6 5.7, respectively) in minimal medium with glycerol as a single carbon source with or without the addition of the lipase inhibitor tetrahydrolipstatin (WT or +THL, respectively). Addition of THL blocks the ability of Mtb to undergo lipid remodeling. D) Addition of THL increases *prpC* expression at pH 7.0 and pH 5.7. *Significantly induced (*p*<0.05) at pH 5.7 relative to pH 7.0 (t test). E) *prpC* is not induced at pH 5.7 with lactate as a single carbon source. Fold change is relative to Mtb grown in glycerol as a sole carbon source at pH 7.0.

Given that endogenous lipid remodeling does not appear to be the source of *prpC* induction at acidic pH, we sought to better understand the conditions leading to *prpCD* induction. Interestingly, although Mtb arrests growth at pH 5.7 with lactate as a single carbon source(10), induction of *prpCD* is not observed in Mtb cultured with lactate as a single carbon source (Figure 3E). This observation suggests that *prpCD* induction during pH 5.7 growth arrest is glycerol-dependent rather than growth arrest dependent. The observation that *prpCD* induction at pH 5.7 does not appear to be required for Mtb growth arrest is consistent with the previous finding that the role of *icl1/2* in growth regulation appears to be independent of the methylcitrate cycle (11).

## Discussion

Transcriptional profiling experiments first indicated that *prpCD* were induced at acidic pH, but only in the presence of glycerol and not pyruvate. Under conditions of robust growth (acidic pH with pyruvate as a carbon source), *prpCD* is slightly repressed at pH 5.7 as compared to pH 7.0. The loss of *prpCD* induction in the presence of vitamin B12 is strongly suggestive that this response is dependent on enhanced accumulation of propionyl-CoA. Interestingly, another cistron that showed the same expression pattern (induced in glycerol and repressed in pyruvate at pH 5.7), includes the Rv2557 and Rv2558 genes. These genes have been previously shown to be induced during starvation and in human granulomas (20). Although, it has previously been shown that Mtb uptakes glycerol during acid growth arrest(11), it is possible that due to a metabolic adaptation that it cannot efficiently utilize glycerol for growth, and thus acid growth arrest, may have some similarities to a carbon starvation response. This would be consistent with the observed TAG catabolism, where it could be utilized as an energy source.

Our initial hypothesis was that *prpCD* is induced during acid growth arrest due to the production of propionyl-CoA generated during lipid remodeling. Lipid remodeling was observed during acid growth arrest, with TAG being catabolized and TDM and SL being synthesized. However, inhibiting lipases with THL resulted in further induction of *prpCD*, a finding contrary to the initial hypothesis. Rather, this finding suggests that lipid remodeling at acidic pH may instead contribute to detoxification of propionyl-CoA, by incorporating the metabolite into methyl-branched cell envelope lipids. The metabolic coupling of propionyl-CoA metabolism to lipid synthesis at acidic pH is consistent with previous work showing lipid synthesis as a readily usable sink for propionyl-CoA incorporation (16, 21, 22). Unlike in this previous work, during acidic pH growth arrest in glycerol-containing medium, the source of propionyl-CoA appears to be endogenous rather than exogenously supplied.

The induction of *prpCD* during acid growth arrest was shown to be dependent on glycerol being present in the media, suggesting that the induction of *prpCD* is dependent on metabolism of glycerol at acidic pH (Figure 4A). To our knowledge, glycerol metabolism leading to the production of propionyl-CoA has not been documented in Mtb. However, in other bacterial species, three separate pathways for the production of propionyl-CoA have been described (23). One of these pathways, the propanediol pathway, involves the conversion of methylgloxal to propionate (24). Given that methylglyoxal can accumulate in Mtb as a byproduct of dihydroxyacetone phosphate (DHAP) metabolism, particularly after inhibition of glycolysis (3), we speculate that *prpCD* induction at acidic pH could be secondary to production of propionyl-CoA via this propanediol pathway or a related pathway (Figure 4B). However, whether Mtb contains enzymes capable of performing this metabolism is not known. Endogenous production of propionyl-CoA does not appear to be a necessary component of acidic pH growth arrest as growth arrested Mtb cultured in minimal medium with lactate does not induce *prpCD* at acidic pH; however, understanding this response could uncover additional aspects of Mtb metabolic remodeling that occurs during acidic pH growth arrest.

**Figure 4.**
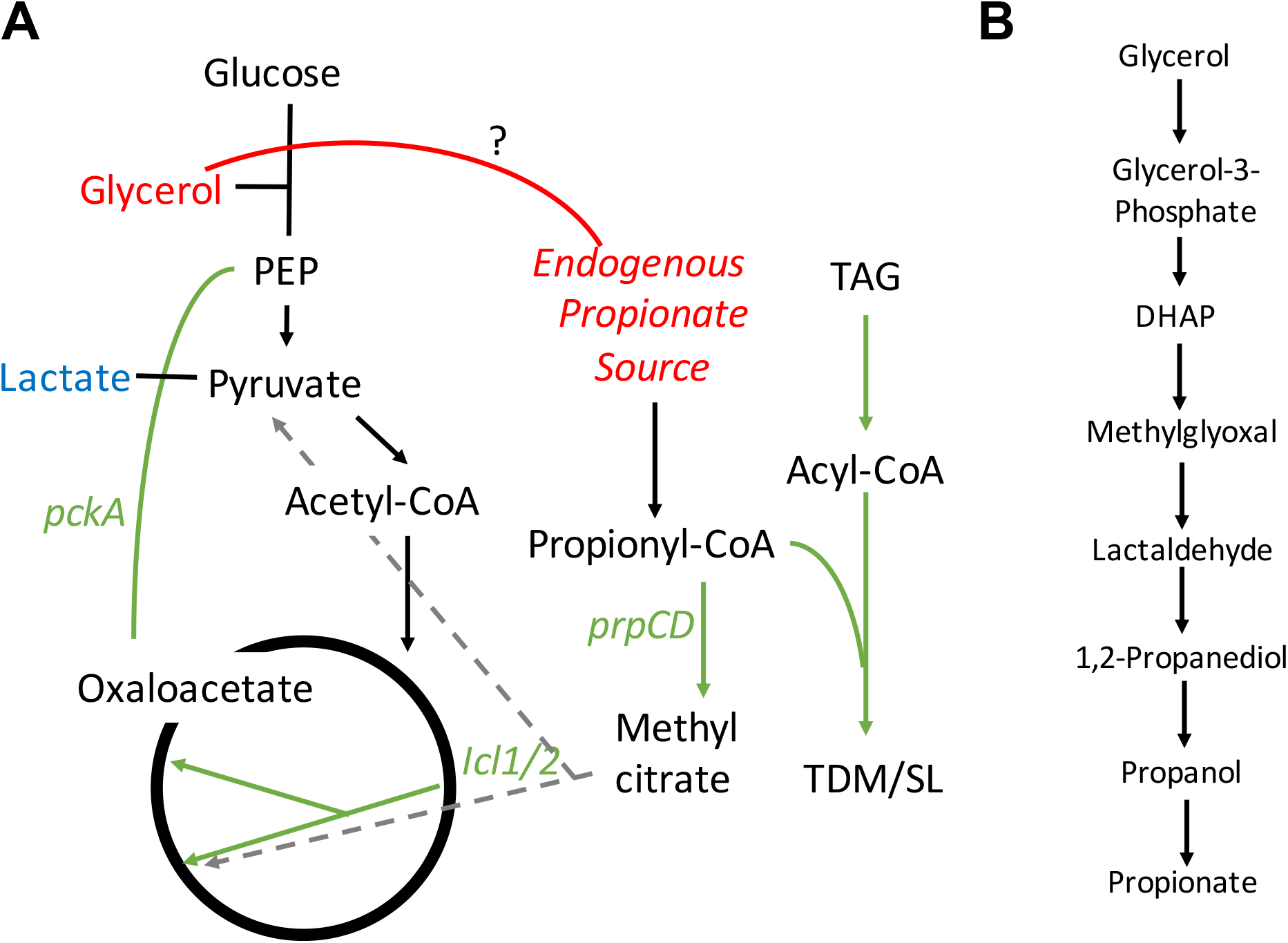
Models for *prpCD* induction and metabolism remodeling during acidic pH growth arrest. A) *prpCD* is induced during acidic pH growth arrest when glycerol is present in the medium, but not when lactate is the single carbon source. It is speculated that the metabolism of glycerol may lead to *de novo* synthesis of propionyl-CoA. The increased expression of *prpCD* in the absence of Mtb lipid remodeling of triacyglycerol (TAG) to sulfolipid (SL) and trehalose dimycolate (TDM) suggests that Mtb uses lipid remodeling as a sink for propionyl-CoA. B) Speculative metabolic pathway for the generation of propionate from glycerol that has been observed in microbes found in the human gut (23).

## Methods

### Bacterial strains and growth conditions

Bacterial growth and strains are as previously described(11). Briefly, all Mtb experiments, unless otherwise stated, were performed with Mtb strain CDC1551. Cultures were maintained in 7H9 Middlebrook medium supplemented with 10% OADC and 0.05% Tween-80 and incubated with 5% CO_2_. All single carbon source experiments were performed in MMAT defined minimal medium as described by Lee *et al*. (22): 1 g/L KH_2_PO_4_, 2.5 g/L Na_2_PO_4_, 0.5 g/L (NH4)_2_SO_4_, 0.15 g/L asparagine, 10 mg/L MgSO_4_, 50 mg/L ferric ammonium citrate, 0.1 mg/L ZnSO_4_, 0.5 mg/L CaCl_2_, and 0.05% Tyloxapol. Medium was buffered using 100 mM MOPS (pH 6.6–7.0) or MES (pH 5.7– 6.5) (25).

### RNA extraction and real time PCR

Mtb cultures were grown at 37°C in T-25 vented, standing tissue culture flasks in 8 mL of a defined minimal medium seeded at an initial OD_600_ of 0.25. After three days, total bacterial RNA was stabilized and extracted as previously described (4). Semi-quantitative real-time PCR was performed using previously described methods (5). Vitamin B12 was supplemented at 10 μg/mL and tetrahydrolipostatin (THL) was added at a concentration of 20 μM. All experiments were conducted with at least two biological replicates and repeated at least twice.

### Analysis of mycobacterial lipids

For lipid remodeling experiments, bacterial cultures were grown in 7H9 +10% OADC with either 8 μCi of [1,2 ^14^C] sodium acetate or [1-^14^C] sodium propionate. Following 15 days of labeling, the bacteria were pelleted and resuspended in the minimal medium containing glycerol as a single carbon source buffered to either pH 7.0 or pH 5.7. At day 0, 6, and 12, two 1 mL aliquots were pelleted and fixed in 4% paraformaldehyde, and the remaining bacteria were pelleted, washed, and the lipids extracted as described previously (10). Total radioactivity and ^14^C incorporation were determined by scintillation counting of the fixed samples and the total extractable lipids, respectively. To analyze lipid species, 5,000 counts per minute (CPM) of the lipid sample was spotted at the origin of 100 cm^2^ silica gel 60 aluminum sheets. To separate sulfolipid for quantification, the TLC was developed with a chloroform:methanol:water (90:10:1 v/v/v) solvent system (18). To separate TAG for quantification, the TLC was developed with a hexane:diethyl ether:acetic acid (80:20:1, v/v/v) solvent system (5). To examine TDM and TMM accumulation the TLC was developed in a chloroform:methanol:ammonium hydroxide (80:20:2 v/v/v) solvent system. Radiolabeled lipids were detected and quantified using a phosphor screen and a Typhoon Imager, and band density quantified using ImageQuant software (26). Radiolabeling experiments, lipid extractions and TLCs were repeated in at least two independent biological replicates with similar findings in both replicates.

## Acknowledgements

We thank Prof. John McKinney for the generous gift of the Δ*icl1/2* mutant and matched wild type control strain used in this study. JB was supported by a Robert J. Schultz student research award. This study was supported by funding from the Michigan State University start-up funds, AgBioResearch and grants from the NIH-NIAID (U54AI057153 and R01AI116605).

## Author Contributions

JB conducted the experiments and JB and RBA designed the experiments, analyzed the data and wrote the manuscript.

